# MAPK13 stabilization via m^6^A modification limits anti-cancer efficacy of rapamycin

**DOI:** 10.1101/2022.08.05.502726

**Authors:** Joohwan Kim, Yujin Chun, Cuauhtemoc B. Ramirez, Lauren A. Hoffner, Sunhee Jung, Ki-Hong Jang, Varvara I. Rubtsova, Cholsoon Jang, Gina Lee

## Abstract

*N*^6^-adenosine methylation (m^6^A) is the most abundant mRNA modification that controls gene expression through diverse mechanisms. m^6^A-dependent regulation of oncogenes and tumor suppressors indeed contribute to tumor development. However, the role of m^6^A-mediated gene regulation after drug treatment or resistance is poorly understood. Here, we report that m^6^A modification of *mitogen-activated protein kinase 13* (*MAPK13*) determines the sensitivity of cancer cells to the mechanistic target of rapamycin complex 1 (mTORC1)- targeting chemotherapeutic agent rapamycin. mTORC1 induces m^6^A modification of *MAPK13* mRNA at its 3’ untranslated region (3’UTR) through methyltransferase-like 3 (METTL3)-METTL14-Wilms’ tumor 1-associating protein (WTAP) methyltransferase complex, thereby stimulating its mRNA degradation via an m^6^A reader protein YTH domain family protein 2 (YTHDF2). Rapamycin blunts this process and stabilizes *MAPK13*. Unexpectedly, *MAPK13* silencing suppresses cell growth and enhances rapamycin’s anti-cancer effects, suggesting that *MAPK13* is an oncogenic gene activated by rapamycin via a feedback regulation. Together, our data indicate that rapamycin-mediated *MAPK13* mRNA stabilization may confer drug resistance, and it can thus be a therapeutic target to sensitize cancer cells to rapamycin.

## Introduction

Transcriptional regulation is a central mechanism required for controlling gene expression. In addition to this canonical process, cells modify genetic material with various chemical moieties as an additional layer of gene regulation. While epigenetic modifications of DNA and histones are well-established, chemical modifications of RNA itself (i.e., epitranscriptomic regulation) have been recently shown to also play crucial roles in gene regulation. Of the RNA modifications, m^6^A is the most abundant (1). m^6^A is deposited on mRNA by a methyltransferase complex, which is composed of three core proteins: METTL3, METTL14, and WTAP (2, 3). m^6^A is mostly enriched on the last exon of mRNA near the stop codon and 3’UTR as revealed by transcriptome-wide sequencing (4, 5). These m^6^A-modified mRNAs then recruit m^6^A-binding “reader” proteins that will determine the diverse fates of these mRNAs. For example, YTHDF destabilizes or promotes the translation of m^6^A-containing transcripts (6, 7).

m^6^A-dependent gene regulation is involved in various biological processes such as stem cell differentiation, embryo development, sex determination, and circadian rhythm; dysregulation of this process can cause various diseases including cancers (8, 9, 10). Interestingly, both increased and decreased m^6^A levels can lead to cancer development, depending on the downstream target genes. METTL3 overexpression in leukemia cells induces the expression of oncogenes such as *cMyc* and *Bcl2* (11). On the other hand, METTL3 downregulation in endometrial cancer induces pro-survival Akt signaling by decreasing the expression of the Akt inhibitor, PHLPP2 (12). Therefore, a comprehensive examination of m^6^A target genes is necessary to better understand the impact of m^6^A modification in different biological contexts.

mTORC1 is overactivated in most human cancers as a master regulator of cell growth (13, 14, 15, 16). The mTORC1 inhibitor rapamycin faces several clinical challenges as chemotherapy, largely due to the developed resistance by tumors or regrowth of tumors after treatment (17, 18, 19). It has been suggested that mTORC1-dependent post-translational modifications of proteins underlie observed rapamycin resistance mechanisms (19). However, whether post-transcriptional RNA modifications confer rapamycin resistance is unknown.

Recent works from our and other labs revealed that activation of m^6^A processing by mTORC1 contributes to tumor progression. mTORC1 increases the expression of METTL3, METTL14, and WTAP, which downregulates cell growth-suppressing genes such as autophagy genes and cMyc suppressor (20, 21, 22). From our transcriptome-wide m^6^A sequencing, we identified additional genes that are potentially regulated by mTORC1-dependent m^6^A modification (20). In this study, we report that a MAPK/p38 isoform, MAPK13/p38d, is a downstream target of the mTORC1-m^6^A-pathway, which likely contributes to the limiting tumor suppressive effects of rapamycin.

## Results

### Identification of genes regulated by mTORC1 and m^6^A writer complex

We previously performed miCLIP-seq in human embryonic kidney HEK293E cells, identifying the genes whose m^6^A level is decreased when total mRNA expression is increased by the mTOR catalytic inhibitor, torin1 (20). Since torin1 suppresses both mTORC1 and mTORC2, we used rapamycin to selectively block mTORC1 and performed a qPCR as a secondary screen (Fig. 1*A*). In parallel, we depleted m^6^A writer complex proteins, METTL3/14 or WTAP, to identify genes that are regulated by m^6^A modification. For these screens, we used lymphangioleiomyomatosis (LAM) patient-derived kidney angiomyolipoma cell line (LAM 621-101), which has an overactivated mTORC1 activity due to TSC2 deficiency (23, 24). Consistent with our previous findings, rapamycin reduced the protein levels of m^6^A writer proteins METTL3, METTL14, and WTAP (Fig. 1, *B* and *C*) (20, 21). We found 10 genes (*BEX1, EIF4A2, EIF6, FGFR3, MAPK13, NOP56, PKD1, SLC25A37, STAT5B*, and *TRP*) whose mRNA levels were induced by rapamycin (Fig. 1*D*). *METTL3/14* knockdown induced mRNA levels of *BEX1, EIF6, MAPK13*, and *SLC25A37* (Fig. 1*E*), and *WTAP* knockdown increased mRNA levels of *EIF6* and *MAPK13* (Fig. 1*F*). Thus, we decided to further study *MAPK13* because its mRNA level was most dramatically and consistently induced by all three conditions (Fig. 1*D-F*).

**Figure 1.**
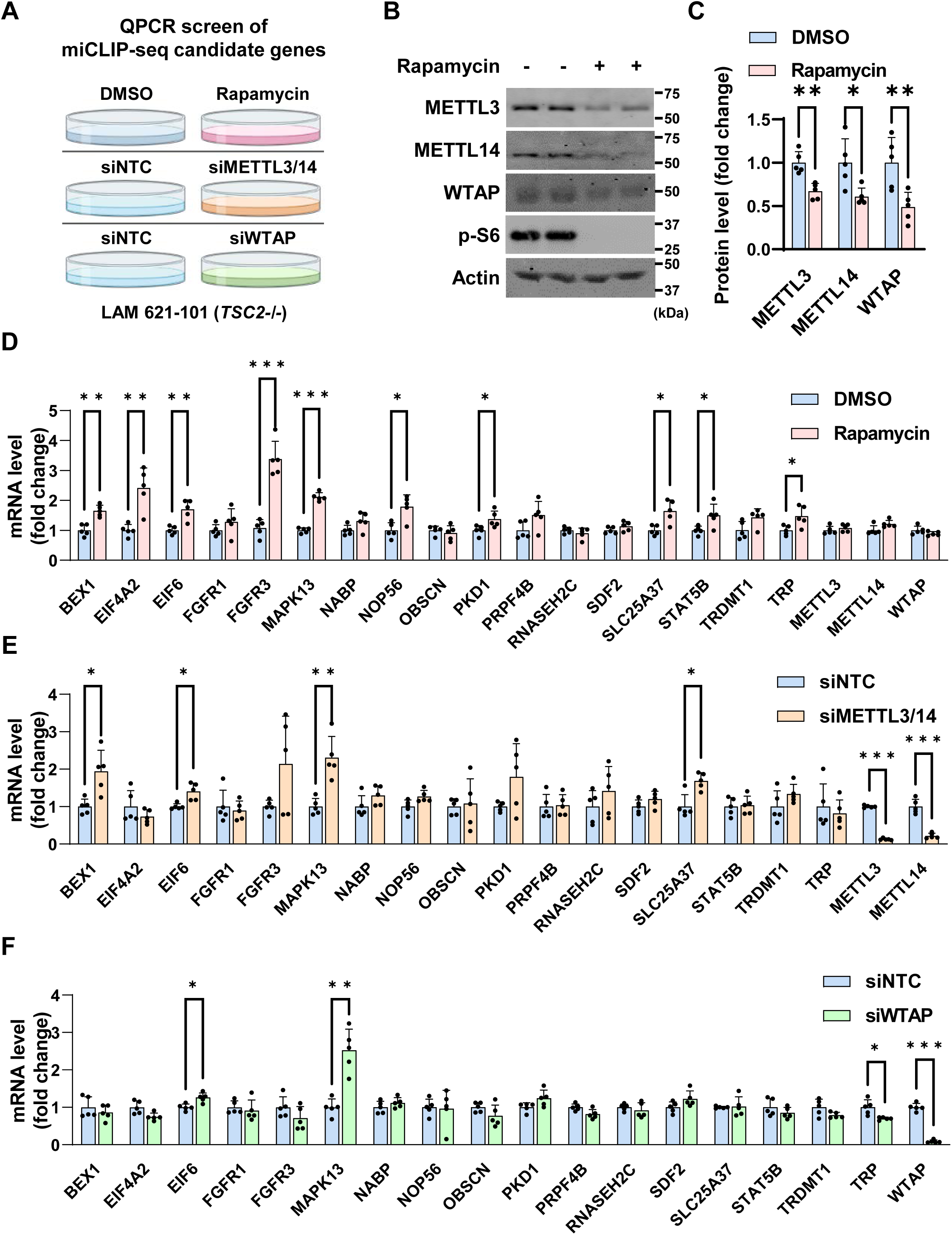
Identification of *MAPK13* as the downstream target of rapamycin and m^6^A writer complex. (A) Schematic of the qPCR screen in LAM 621-101 (*TSC2-/-*) cells to identify target genes whose expression is regulated by rapamycin and m^6^A writer complex. The screen sets include three conditions treated with DMSO (control) vs. rapamycin for 48 h, transfected with siNTC (control) vs. siMETTL3/14, and transfected with siNTC vs. siWTAP. Candidate genes were selected from our previous miCLIP-seq in HEK293E cells treated with mTOR inhibitor, torin1 (20). (B, C) Immunoblot analysis of LAM 621-101 cells treated with DMSO or rapamycin. (C) shows a quantification graph of immunoblot bands. N=5. (D-F) qPCR analysis of 17 candidate genes in LAM 621-101 cells treated with DMSO vs. rapamycin (D), transfected with siNTC vs. siMETTL3/14 (E), or transfected with siNTC vs. siWTAP (F). N=5. *p < 0.05, **p < 0.01, ***p < 0.001. Error bars show standard deviation (SD). Numbers on the immunoblot indicate the positions of molecular weight markers.

### m^6^A modification regulates MAPK13 expression among p38 isoforms

Next, we assessed protein levels of MAPK13 to examine whether the changes in *MAPK13* mRNA levels are reflected in MAPK13 protein expression. Upon rapamycin treatment, the protein levels of MAPK13 increased by 2-fold (Fig. 2, *A* and *B*). Double knockdown of *METTL3/14* also resulted in a 2-fold increase in MAPK13 protein expression (Fig. 2, C and *D*). Overall, the extent of protein induction (Fig. 2, *A-D*) matched well with the increases in mRNA levels (Fig. 1, *D-F*).

**Figure 2.**
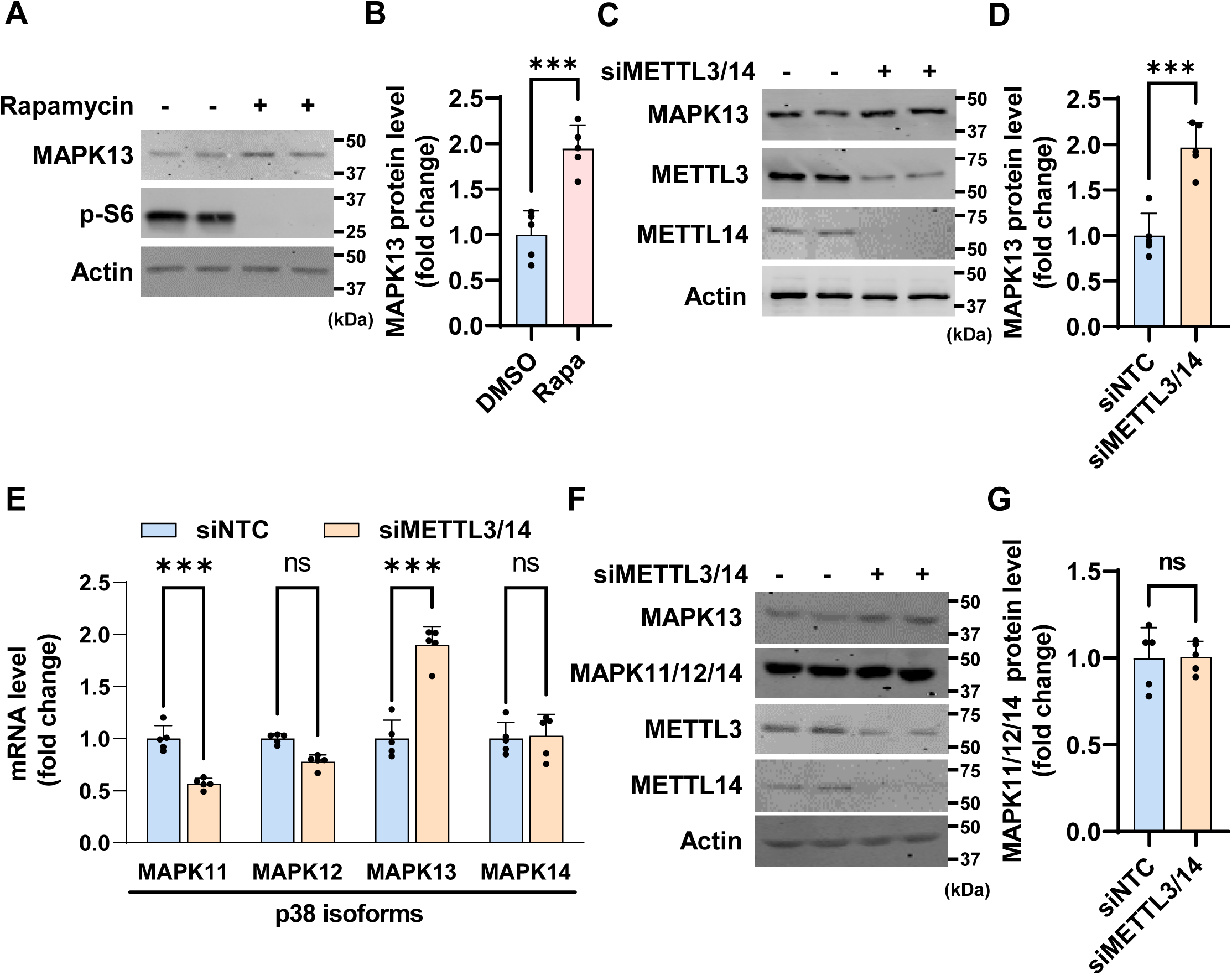
mTORC1 and m^6^A regulate MAPK13 expression among the other p38 MAPK isoforms. (A, B) Immunoblot analysis of LAM 621-101 cells treated with DMSO or rapamycin. (B) shows the quantification graph of immunoblot bands. N=5. (C, D) Immunoblot analysis of LAM 621-101 cells transfected with siNTC or siMETTL3/14. (D) shows a quantification graph of immunoblot bands. N=5. (E) qPCR analysis of p38 MAPK family genes. LAM 621-101 cells were transfected with siNTC or siMETTL3/14. N=5. (F, G) Immunoblot analysis of LAM 621-101 cells transfected with siNTC or siMETTL3/14. (G) shows a quantification graph of immunoblot bands. N=5. ***p < 0.001, ns = not significant. Error bars show standard deviation (SD). Numbers on the immunoblot indicate the positions of molecular weight markers.

MAPK13 is a member of the p38 MAPK protein family composed of p38α (MAPK14), p38β (MAPK11), p38γ (MAPK12), and p38d (MAPK13). These proteins control diverse cellular signaling processes, including proliferation, differentiation, inflammation, and cell death responses (25, 26, 27, 28). Interestingly, in contrast to *MAPK13*, knockdown of *METTL3/14* did not induce mRNA levels of *MAPK11, MAPK12*, or *MAPK14* (Fig. 2*E*). In the case of *MAPK11*, its mRNA level was decreased (Fig. 2*E*). To examine the protein level of these isoforms, we used an antibody that detects amino acid sequences across three p38 isoforms, MAPK11, MAPK12, and MAPK14 (29). This antibody does not detect MAPK13. Interestingly, the protein expression of these p38 isoforms (MAPK11, MAPK12, and MAPK14) did not change regardless of *METTL3/14* knockdown (Fig. 2, *F* and *G*). Thus, MAPK13 is a unique p38 isoform regulated by mTORC1-dependent m^6^A modification.

### mTORC1-m^6^A-YTHDF2 destabilizes *MAPK13* mRNA

Once modified with m^6^A, mRNAs recruit m^6^A reader proteins that determine the fate of target transcripts such as mRNA stability or translation efficiency (6, 7). From our previous miCLIP-seq, we found that a specific m^6^A modification site on 3’UTR of *MAPK13* was decreased by mTOR inhibition (Fig. 3*A*) (20). Given that suppression of such m^6^A modification by *METTL3/14* or *WTAP* knockdown increases both MAPK13 mRNA and protein levels (Fig. 1 and 2), we hypothesized that *MAPK13* mRNA is degraded by YTHDF2, an m^6^A reader protein that destabilizes target transcripts (6, 7, 30, 31). Consistent with our hypothesis, knockdown of *YTHDF2* resulted in a significant increase in *MAPK13* mRNA levels (Fig. 3*B*). Consequently, MAPK13 protein levels also increased (Fig. 3*C*). Hence, YTHDF2 is the effector protein responsible for *MAPK13* mRNA degradation after mTORC1-mediated m^6^A modification.

**Figure 3.**
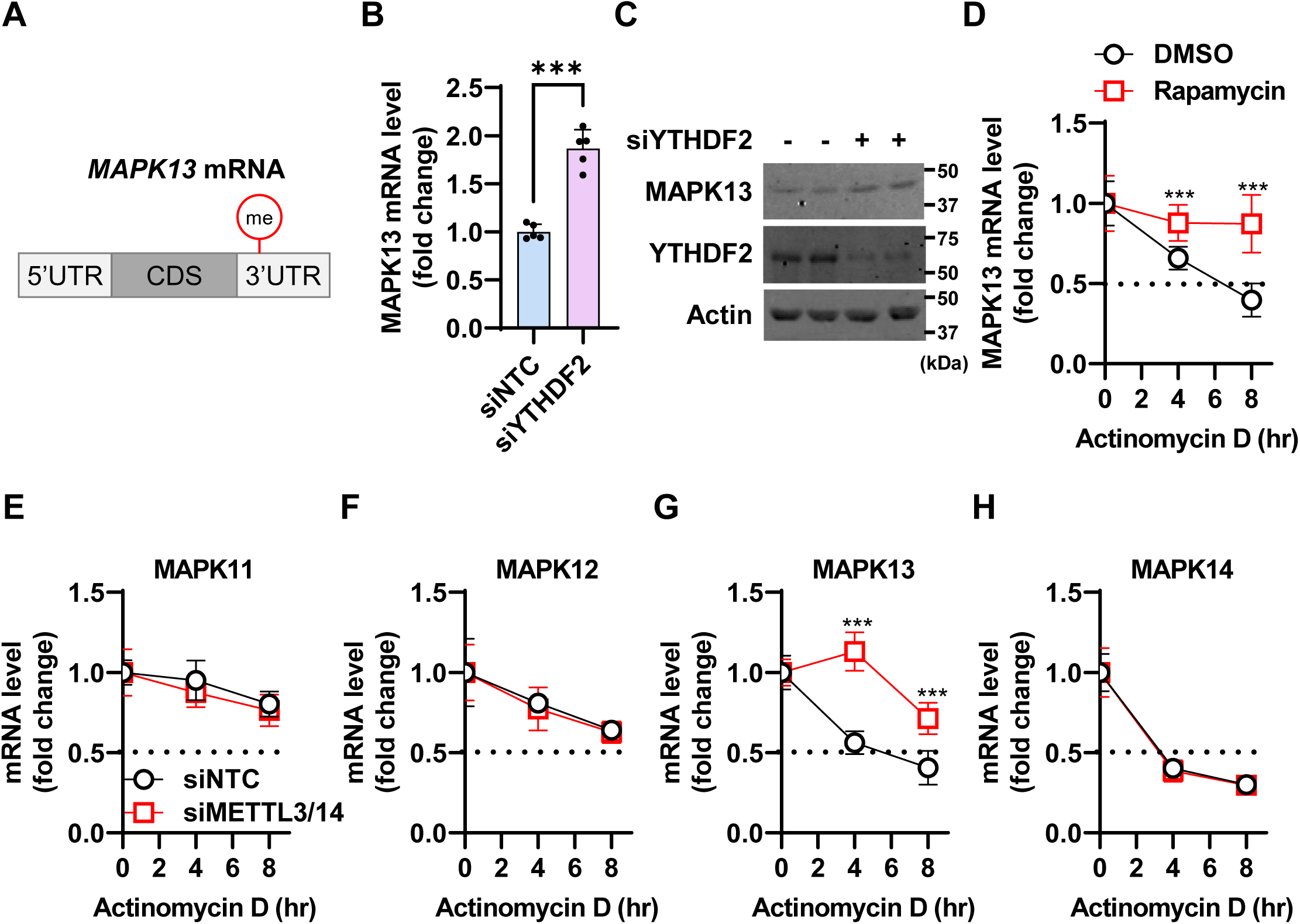
*MAPK13* mRNA stability is regulated by the mTORC1-m^6^A-YTHDF2 axis. (A) Schematic of *MAPK13* mRNA. From miCLIP-seq (Cho et al., 2021) (20), an mTORC1-dependent m^6^A modification site (red) was identified on the 3’UTR of *MAPK13*. (B, C) qPCR and immunoblot analysis of LAM 621-101 cells transfected with siNTC or siYTHDF2. N=5. (D) mRNA stability analysis of *MAPK13* in LAM 621-101 cells treated with rapamycin. Cells were treated with actinomycin D for the indicated times. N=5. (E-H) mRNA stability analysis of *MAPK11* (E), *MAPK12* (F), *MAPK13* (G), and *MAPK14* (H) in LAM 621-101 cells transfected with siNTC or siMETTL3/14. Cells were treated with actinomycin D for the indicated times. N=5. ***p < 0.001. Error bars show standard deviation (SD). Numbers on the immunoblot indicate the positions of molecular weight markers.

To further confirm whether the stability of *MAPK13* mRNA is indeed regulated by mTORC1-dependent m^6^A modification, we sought to quantitate mRNA half-life. To this end, we treated actinomycin D to block *de novo* mRNA synthesis and measured the remaining transcript levels at different time points (32). In control cells, *MAPK13* mRNA was degraded in a time-dependent manner with a half-life of 6 h (Fig. 3*D*). However, upon rapamycin treatment, the stability of *MAPK13* mRNA was dramatically increased, with 90% of transcripts remaining even after 8 h (Fig. 3*D*). Similarly, *METTL3/14* double knockdown also markedly increased the half-life of *MAPK13* mRNA but not that of the other p38 isoforms (Fig. 3, *E-H*). Collectively, these results demonstrate that rapamycin increases the mRNA stability of *MAPK13* via the m^6^A-YTHDF2 axis.

### MAPK13 inhibition enhances rapamycin’s anti-tumor effects

Among various mitogen-activated MAPK family member proteins, p38 MAPKs are mostly responsible for reduced cell growth or increased apoptosis (25, 26, 27, 28). We therefore reasoned that mTORC1-activated cancer cells actively degrade *MAPK13* mRNA to promote their cell growth. To test this hypothesis, we knocked down *MAPK13* in LAM cells and measured cell proliferation (Fig. 4*A*). However, to our surprise, *MAPK13* knockdown preferentially reduced cell growth, indicating that MAPK13 plays a role in promoting cell growth. In the absence of MAPK13, rapamycin treatment further suppressed cell growth (Fig. 4*B*). *MAPK13* knockdown also augmented cell migration suppression by rapamycin (Fig. 4, *C* and *D*). Thus, MAPK13 induction by rapamycin treatment may limit the tumor suppressive effects of the drug. These findings also suggest that inhibition of MAPK13 coupled with rapamycin can be more effective at impairing tumor growth compared to sole rapamycin treatment (Fig. 4*E*).

**Figure 4.**
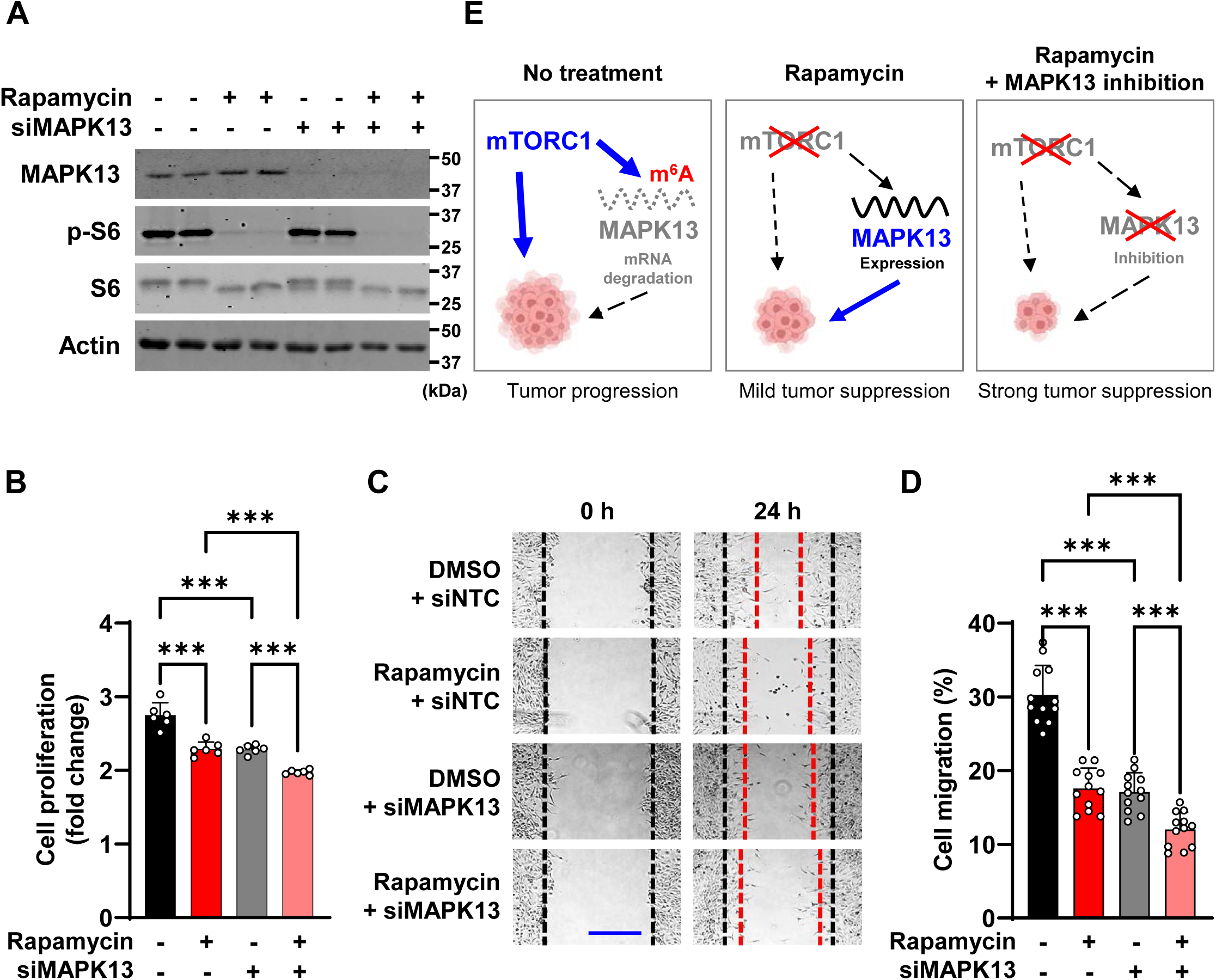
MAPK13 inhibition enhances rapamycin’s suppressive effect on cancer cell growth and migration. (A) Immunoblot analysis of LAM 621-101 cells transfected with siNTC or siMAPK13, followed by treatment with DMSO or rapamycin. N=5. (B) Cell proliferation assay of LAM 621-101 cells transfected with siNTC or siMAPK13, followed by treatment with DMSO or rapamycin. The graph shows the fold increase of cell numbers 3 days after the treatment. N=6. (C, D) Wound healing assay of LAM 621-101 cells transfected with siNTC or siMAPK13, followed by treatment with DMSO or rapamycin. After scratching the cell layer to form a wound, images were captured at 0 h and 24 h to evaluate cell migration. Black dotted lines indicate the initial wound area; red dotted lines mark the migrating front of cells (C). Cell migration efficiency was calculated by measuring the wound area at each time point by Image J software (D). Scale bar, 500 µm. N=12. (E) A schematic diagram describing the regulation of *MAPK13* expression by mTORC1-dependent m^6^A methylation (left, without rapamycin; middle, with rapamycin) and the potential impact of MAPK13 inhibition in tumor suppression in combination with rapamycin treatment (right). ***p < 0.001. Error bars show standard deviation (SD). Numbers on the immunoblot indicate the positions of molecular weight markers.

## Discussion

Because of its specific inhibitory activity on mTORC1, rapamycin was thought to be a ground-breaking anti-cancer therapeutic for a broad spectrum of mTORC1-overactivated human cancers. However, clinical trials revealed that rapamycin was not as efficient as expected. Some tumors even regrow into a bigger size after cessation of rapamycin treatment (17, 18, 19). One of the mechanisms for rapamycin resistance is activation of other growth-promoting signaling pathways (19). In breast cancer patients, mitogenic ERK MAPK signaling was increased in cancer tissues upon rapamycin treatment (33). This unexpected observation led to the identification of negative feedback signaling pathways downstream of mTORC1; while mTORC1 promotes anabolic pathways for cell growth, it ironically inhibits several pro-growth signals including PI3K, Ras, and MEK (19). Some of these pro-growth signals such as PI3K and RAS are upstream activators of mTORC1; therefore, when mTORC1 is suppressed by rapamycin, these negative feedbacks are released, resulting in the continued growth of cancer cells (34). On the other hand, co-treatment rapamycin with PI3K or MEK inhibitors is more effective for tumor suppression in cell culture and mouse models (33, 35). Here, we identified MAPK13 as another player that potentially limits rapamycin’s tumor suppressive effects. MAPK13 has been shown to activate mTORC1, indicating the negative feedback loop between MAPK13 and mTORC1 (36). Indeed, suppression of MAPK13 in combination with rapamycin treatment enhanced rapamycin’s effect on cell growth and migration inhibition, suggesting MAPK13 as a potential therapeutic target for augmenting rapamycin sensitivity (Fig. 4).

MAPK13 is one of the four p38 MAPK family proteins. Among the isoforms, MAPK14/p38α is expressed in most cell types, whereas the other MAPK family genes are expressed in specific tissues; MAPK11/p38β and MAPK12/p38γ are expressed in the brain and skeletal muscle, while MAPK13/p38d is expressed in the kidney and lung (28). Intriguingly, the kidney and lung are the two dominant organs that develop tumors in LAM patients (37). Our data indicate that, among the p38 MAPK family genes, only MAPK13/p38d was regulated by mTORC1-dependent m^6^A modification (Fig. 2 and 3). Therefore, small molecule inhibitors that specifically target MAPK13/p38d isoform can be a selective therapeutic regimen with improved efficacy and lower toxicity. While MAPK14/p38α isoform has been most extensively studied, MAPK13/p38d has recently emerged as a potential drug target because of its roles in stress responses, cytokine production, and tumor development (27, 26, 38, 39). In this sense, our findings suggest MAPK13 as a promising target for combination chemotherapy with rapamycin to overcome the limited efficacy of rapamycin.

## Experimental Procedures

### Cell culture

Human kidney angiomyolipoma-derived LAM 621-101 cell line (mTORC1 overactivation due to TSC2 deficiency; RRID: CVCL_S879) was provided by Drs. Jane Yu and Elisabeth Henske (23). LAM 621-101 cells were grown in DMEM (GIBCO) with 10% FBS (Sigma-Aldrich) at 37°C with 5% CO2. 5×10^6 cells counted by Multisizer 4e Coulter Counter (Beckman) were plated on a 60 mm plate and serum starved before analysis unless otherwise indicated. Rapamycin (Calbiochem) dissolved in DMSO was treated at the final concentration of 20 nM. siRNAs (Sigma-Aldrich) dissolved in nuclease-free water were transfected using Lipofectamine RNAiMAX reagent (Invitrogen) at the final concentration of 30 nM. siRNA list is provided in Table S1.

### Quantitative PCR (qPCR)

PureLink RNA isolation kit (Life Technologies) was used to isolate total RNA from cells. After removing genomic DNA by DNase I (Sigma-Aldrich-Aldrich), RNA was reverse transcribed to cDNA using the iScript kit (Bio-rad). The resulting cDNA was analyzed by quantitative RT-PCR (qPCR) using SYBR green master mix (Life Technologies) on QuantStudio6 Real-Time PCR system (Life Technologies). For the qPCR screen in Figure 1, 17 final candidate genes from our previous miCLIP-seq performed in HEK293E cells with and without mTOR inhibitor, torin1, were used (Cho et al., 2021) (20). mRNA levels were calculated by delta-delta CT method using housekeeping genes *Actin, PPIB*, and *TBP*. The primer list is provided in Table S2.

### Immunoblot

Cells were homogenized on ice using RIPA lysis buffer (25 mM Tris-HCl [pH 7.4], 2 mM EDTA, 150 mM NaCl, 0.1% SDS, 2 mM DTT, 1% Sodium deoxycholate, 1% NP-40) supplemented with protease inhibitors (1 mM PMSF, 2 μg/mL pepstatin A, 10 μg/mL leupeptin, 10 μg/mL aprotinin) and phosphatase inhibitors (10 mM NaF, 1 mM Na3VO4). Cell lysates were cleared by centrifugation at 13,000 rpm at 4°C for 30 min. Detergent compatible (DC) protein assay (Bio-rad) was used to measure protein concentration. Proteins were boiled for 10 min with Laemmli sample buffer. SDS-PAGE gels were used to separate proteins (10-30 μg) and transferred to the nitrocellulose membrane (Amersham Biosciences). Membranes were then incubated with Odyssey blocking solution (Li-COR Biosciences), followed by incubation with primary and IRDye secondary (Li-COR Biosciences) antibodies. Immunoblot signals were detected and quantified by Image Studio software with the Li-COR imaging system (Li-COR Biosciences). Immunoblot images are representative of at least two independent experiments. Antibodies against pS6(S240/S244), S6, Beta-actin, p38 (MAPK11/12/14), METTL3, and METTL14 (Cell Signaling Technology); YTHDF2 (Proteintech); WTAP (Abcam); and MAPK13 (R&D Systems) were used.

### mRNA stability assay

Cells were treated with 5 μg/ml actinomycin D (Sigma-Aldrich) to inhibit transcription and were collected at 0, 4, and 8 h to analyze the remaining mRNA levels. Total RNA was extracted and mRNA levels were analyzed by qPCR as described above.

### Cell proliferation assay

siRNA-transfected cells were seeded on a 60 mm plate. After 24 h, cells were treated with DMSO (control) or rapamycin without FBS. Cell numbers were measured using Multisizer 4e Coulter Counter (Beckman) at 0 h and 72 h after treatment. Cell proliferation (fold change) was calculated by dividing the cell numbers at 72 h by the cell numbers at 0 h.

### Cell migration assay

Wound-healing assay was applied to assess cell migration. siRNA-transfected cells were seeded on a 6-well plate. After 24 h, cells were treated with DMSO (control) or rapamycin without FBS. Once cells are confluent, a clear wound line was created using a sterile 200 μL pipette tip. Cell images containing the wound area were taken at 0 h and 24 h using Eclipse Ts2-FL microscope and DS-Fi3 Camera (Nikon). Cell migration efficiency (%) was calculated by measuring the cell migration area (0 h ∼ 24 h) using the ImageJ software program.

### Statistical analysis

Statistical analyses were performed using GraphPad Prism software. All values are presented as mean ± standard deviation (SD). Statistical significance was determined using a two-tailed Student’s t-test for comparison between two groups or multiple unpaired t-tests for comparison between multiple groups. Statistical significance is presented as *p < 0.05, **p < 0.01, ***p < 0.001, or ns = not significant.

## Abbreviations and nomenclature

Bcl-2: (B-cell lymphoma 2)
PHLPP2: (PH domain and leucine-rich repeat protein phosphatase 2)
LAM: (Lymphangioleiomyomatosis)
TSC: (Tuberous sclerosis complex)
miCLIP-seq: (m6A individual-nucleotide resolution UV crosslinking and immunoprecipitation)
BEX1: (Brain expressed X-linked 1)
EIF4A2: (Eukaryotic translation initiation factor 4A2)
EIF6: (Eukaryotic translation initiation factor 6)
FGFR: (Fibroblast growth factor receptor)
NABP1: (Nucleic acid binding protein 1)
NOP56: (NOP56 ribonucleoprotein)
OBSCN: (Obscurin)
PKD: (Polycystic kidney disease)
PRPF4B: (Pre-mRNA processing factor 4B)
SDF2: (Stromal cell-derived factor 2)
SLC25A37: (Solute carrier family 25 member 37)
STAT5B: (Signal transducer and activator of transcription 5B)
TRDMT1: (tRNA aspartic acid methyltransferase 1)
TRP: (Transient receptor potential)
ERK: (Extracellular signal-regulated kinase)
PI3K: (Phosphoinositide 3-kinase).

## Acknowledgment

We are grateful to Drs. Jane Yu and Elisabeth Henske for sharing a patient-derived LAM 621-101 cell line. We also thank members of the Lee and Jang laboratories, and Drs. Yongsheng Shi and Dean Aguiar for technical assistance and scientific discussions. This research was supported by DOD TS200022 (G.L.), NIH K22CA234399 (G.L.), R01AA029124 (C.J.), and P30CA062203 (UCI CFCCC Anti-Cancer Challenge award). S.J. was supported by a postdoctoral fellowship from the National Research Foundation of Korea (2021R1A6A3A14039681). C.B.R. was supported by a predoctoral fellowship from the UCI IMSD program (R25GM055246). Schematics in the Figures are created with BioRender.

## Supplemental Information

**Table S1. siRNA list**.

**Table S2. Primer list**.

## References

1. Desrosiers, R., Friderici, K., and Rottman, F. (1974) Identification of methylated nucleosides in messenger RNA from Novikoff hepatoma cells. Proc. Natl. Acad. Sci. U. S. A. 71, 3971–3975

2. Karthiya, R., and Khandelia, P. (2020) m6A RNA Methylation: Ramifications for Gene Expression and Human Health. Mol. Biotechnol. 62, 467–484

3. Huang, H., Weng, H., and Chen, J. (2020) m6A Modification in Coding and Non-coding RNAs: Roles and Therapeutic Implications in Cancer. Cancer Cell. 37, 270–288

4. Dominissini, D., Moshitch-Moshkovitz, S., Schwartz, S., Salmon-Divon, M., Ungar, L., Osenberg, S., Cesarkas, K., Jacob-Hirsch, J., Amariglio, N., Kupiec, M., Sorek, R., and Rechavi, G. (2012) Topology of the human and mouse m6A RNA methylomes revealed by m6A-seq. Nature. 485, 201–206

5. Meyer, K. D., Saletore, Y., Zumbo, P., Elemento, O., Mason, C. E., and Jaffrey, S. R. (2012) Comprehensive analysis of mRNA methylation reveals enrichment in 3′ UTRs and near stop codons. Cell. 149, 1635–1646

6. Zaccara, S., and Jaffrey, S. R. (2020) A Unified Model for the Function of YTHDF Proteins in Regulating m6A-Modified mRNA. Cell. 181, 1582-1595.e18

7. Lee, Y., Choe, J., Park, O. H., and Kim, Y. K. (2020) Molecular Mechanisms Driving mRNA Degradation by m6A Modification. Trends Genet. 36, 177–188

8. Wu, J., Frazier, K., Zhang, J., Gan, Z., Wang, T., and Zhong, X. (2020) Emerging role of m6A RNA methylation in nutritional physiology and metabolism. Obes. Rev. 21, 1–10

9. Wang, T., Kong, S., Tao, M., and Ju, S. (2020) The potential role of RNA N6-methyladenosine in Cancer progression. Mol. Cancer. 19, 1–18

10. Jaffrey, S. R., and Kharas, M. G. (2017) Emerging links between m6A and misregulated mRNA methylation in cancer. Genome Med. 9, 8–10

11. Vu, L. P., Pickering, B. F., Cheng, Y., Zaccara, S., Nguyen, D., Minuesa, G., Chou, T., Chow, A., Saletore, Y., Mackay, M., Schulman, J., Famulare, C., Patel, M., Klimek, V. M., Garrett-Bakelman, F. E., Melnick, A., Carroll, M., Mason, C. E., Jaffrey, S. R., and Kharas, M. G. (2017) The N 6 -methyladenosine (m 6 A)-forming enzyme METTL3 controls myeloid differentiation of normal hematopoietic and leukemia cells. Nat. Med. 23, 1369–1376

12. Jun Liu, Mark A Eckert, Bryan T Harada, Song-Mei Liu, Zhike Lu, Kangkang Yu, Samantha M Tienda3, Agnieszka Chryplewicz3, Allen C Zhu1, 2, 6, Ying Yang4, Jing-Tao Huang4, Shao-Min Chen4, Zhi-Gao Xu7, Xiao-Hua Leng8, Xue-Chen Yu9, Jie Cao10, Zezhou Zhang10, and C. H. (2018) m6A mRNA methylation regulates AKT activity to promote the proliferation. Nat Cell Biol. 176, 139–148

13. Menon, S., and Manning, B. D. (2008) Common corruption of the mTOR signaling network in human tumors. Oncogene. 27, S43–S51

14. Zou, Z., Tao, T., Li, H., and Zhu, X. (2020) MTOR signaling pathway and mTOR inhibitors in cancer: Progress and challenges. Cell Biosci. 10, 1–11

15. Mossmann, D., Park, S., and Hall, M. N. (2018) mTOR signalling and cellular metabolism are mutual determinants in cancer. Nat. Rev. Cancer. 18, 744–757

16. Kim, J., and Guan, K. L. (2019) mTOR as a central hub of nutrient signalling and cell growth. Nat. Cell Biol. 21, 63–71

17. Bissler, J. J., McCormack, F. X., Young, L. R., Elwing, J. M., Chuck, G., Leonard, J. M., Schmithorst, V. J., Laor, T., Brody, A. S., Bean, J., Salisbury, S., and Franz, D. N. (2008) Sirolimus for Angiomyolipoma in Tuberous Sclerosis Complex or Lymphangioleiomyomatosis. N. Engl. J. Med. 358, 140–151

18. Marsh, D. J., Trahair, T. N., Martin, J. L., Chee, W. Y., Walker, J., Kirk, E. P., Baxter, R. C., and Marshall, G. M. (2008) Rapamycin treatment for a child with germline PTEN mutation. Nat. Clin. Pract. Oncol. 5, 357–361

19. Li, J., Kim, S. G., and Blenis, J. (2014) Rapamycin: One drug, many effects. Cell Metab. 19, 373–379

20. Cho, S., Lee, G., Pickering, B. F., Perrimon, N., Jaffrey, S. R., Cho, S., Lee, G., Pickering, B. F., Jang, C., Park, J. H., He, L., and Mathur, L. (2021) Article mTORC1 promotes cell growth via m 6 A-dependent mRNA degradation ll Article mTORC1 promotes cell growth via m 6 A-dependent mRNA degradation. Mol. Cell. 81, 2064-2075.e8

21. Tang, H. W., Weng, J. H., Lee, W. X., Hu, Y., Gu, L., Cho, S., Lee, G., Binari, R., Li, C., Cheng, M. E., Kim, A. R., Xu, J., Shen, Z., Xu, C., Asara, J. M., Blenis, J., and Perrimon, N. (2021) mTORC1-chaperonin CCT signaling regulates m6A RNA methylation to suppress autophagy. Proc. Natl. Acad. Sci. U. S. A. 118, 1–8

22. Villa, E., Sahu, U., O’Hara, B. P., Ali, E. S., Helmin, K. A., Asara, J. M., Gao, P., Singer, B. D., and Ben-Sahra, I. (2021) mTORC1 stimulates cell growth through SAM synthesis and m6A mRNA-dependent control of protein synthesis. Mol. Cell. 81, 2076-2093.e9

23. Yu, J., Astrinidis, A., Howard, S., and Henske, E. P. (2004) Estradiol and tamoxifen stimulate LAM-associated angiomyolipoma cell growth and activate both genomic and nongenomic signaling pathways. Am. J. Physiol. -Lung Cell. Mol. Physiol. 286, 694–700

24. Yu, J. J., Robb, V. A., Morrison, T. A., Ariazi, E. A., Karbowniczek, M., Astrinidis, A., Wang, C., Hernandez-Cuebas, L., Seeholzer, L. F., Nicolas, E., Hensley, H., Jordan, V. C., Walker, C. L., and Henske, E. P. (2009) Estrogen promotes the survival and pulmonary metastasis of tuberin-null cells. Proc. Natl. Acad. Sci. U. S. A. 106, 2635–2640

25. Asih, P. R., Prikas, E., Stefanoska, K., Tan, A. R. P., Ahel, H. I., and Ittner, A. (2020) Functions of p38 MAP Kinases in the Central Nervous System. Front. Mol. Neurosci. 13, 1–27

26. Anton, D. B., Ducati, R. G., Timmers, L. F. S. M., Laufer, S., and Goettert, M. I. (2021) A special view of what was almost forgotten: P38d mapk. Cancers (Basel). 10.3390/cancers13092077

27. Tan, F. L. S., Ooi, A., Huang, D., Wong, J. C., Qian, C. N., Chao, C., Ooi, L., Tan, Y. M., Chung, A., Cheow, P. C., Zhang, Z., Petillo, D., Yang, X. J., and Teh, B. T. (2010) p38delta/MAPK13 as a diagnostic marker for cholangiocarcinoma and its involvement in cell motility and invasion. Int. J. Cancer. 126, 2353–2361

28. Escós, A., Risco, A., Alsina-Beauchamp, D., and Cuenda, A. (2016) p38γ and p38d mitogen activated protein kinases (MAPKs), new stars in the MAPK galaxy. Front. Cell Dev. Biol. 4, 1–7

29. Yang, C., Zhu, Z., Tong, B. C. K., Iyaswamy, A., Xie, W. J., Zhu, Y., Sreenivasmurthy, S. G., Senthilkumar, K., Cheung, K. H., Song, J. X., Zhang, H. J., and Li, M. (2020) A stress response p38 MAP kinase inhibitor SB202190 promoted TFEB/TFE3-dependent autophagy and lysosomal biogenesis independent of p38. Redox Biol. 32, 101445

30. Du, H., Zhao, Y., He, J., Zhang, Y., Xi, H., Liu, M., Ma, J., and Wu, L. (2016) YTHDF2 destabilizes m 6 A-containing RNA through direct recruitment of the CCR4-NOT deadenylase complex. Nat. Commun. 10.1038/ncomms12626

31. Park, O. H., Ha, H., Lee, Y., Boo, S. H., Kwon, D. H., Song, H. K., and Kim, Y. K. (2019) Endoribonucleolytic Cleavage of m6A-Containing RNAs by RNase P/MRP Complex. Mol. Cell. 74, 494-507.e8

32. Shyu, A. B., Greenberg, M. E., and Belasco, J. G. (1989) The c-fos transcript is targeted for rapid decay by two distinct mRNA degradation pathways. Genes Dev. 3, 60–72

33. Carracedo, A., Ma, L., Teruya-Feldstein, J., Rojo, F., Salmena, L., Alimonti, A., Egia, A., Sasaki, A. T., Thomas, G., Kozma, S. C., Papa, A., Nardella, C., Cantley, L. C., Baselga, J., and Pandolfi, P. P. (2008) Inhibition of mTORC1 leads to MAPK pathway activation through a PI3K-dependent feedback loop in human cancer. J. Clin. Invest. 118, 3065–3074

34. Mendoza, M. C., Er, E. E., and Blenis, J. (2011) The Ras-ERK and PI3K-mTOR pathways: Cross-talk and compensation. Trends Biochem. Sci. 36, 320–328

35. Lu, Y., Zhang, E. Y., Liu, J., and Yu, J. J. (2020) Inhibition of the mechanistic target of rapamycin induces cell survival via MAPK in tuberous sclerosis complex. Orphanet J. Rare Dis. 15, 1–11

36. Linares, J. F., Duran, A., Reina-Campos, M., Aza-Blanc, P., Campos, A., Moscat, J., and Diaz-Meco, M. T. (2015) Amino Acid Activation of mTORC1 by a PB1-Domain-Driven Kinase Complex Cascade. Cell Rep. 12, 1339–1352

37. Henske, E. P., Józwiak, S., Kingswood, J. C., Sampson, J. R., and Thiele, E. A. (2016) Tuberous sclerosis complex. Nat. Rev. Dis. Prim. 10.1038/nrdp.2016.35

38. Liu, Y., Chang, Y., and Cai, Y. (2020) circTNFRSF21, a newly identified circular RNA promotes endometrial carcinoma pathogenesis through regulating miR-1227-MAPK13/ATF2 axis. Aging (Albany. NY). 12, 6774–6792

39. Bouhaddou, M., Memon, D., Meyer, B., White, K. M., Rezelj, V. V., Correa Marrero, M., Polacco, B. J., Melnyk, J. E., Ulferts, S., Kaake, R. M., Batra, J., Richards, A. L., Stevenson, E., Gordon, D. E., Rojc, A., Obernier, K., Fabius, J. M., Soucheray, M., Miorin, L., Moreno, E., Koh, C., Tran, Q. D., Hardy, A., Robinot, R., Vallet, T., Nilsson-Payant, B. E., Hernandez-Armenta, C., Dunham, A., Weigang, S., Knerr, J., Modak, M., Quintero, D., Zhou, Y., Dugourd, A., Valdeolivas, A., Patil, T., Li, Q., Hüttenhain, R., Cakir, M., Muralidharan, M., Kim, M., Jang, G., Tutuncuoglu, B., Hiatt, J., Guo, J. Z., Xu, J., Bouhaddou, S., Mathy, C. J. P., Gaulton, A., Manners, E. J., Félix, E., Shi, Y., Goff, M., Lim, J. K., McBride, T., O’Neal, M. C., Cai, Y., Chang, J. C. J., Broadhurst, D. J., Klippsten, S., De wit, E., Leach, A. R., Kortemme, T., Shoichet, B., Ott, M., Saez-Rodriguez, J., tenOever, B. R., Mullins, R. D., Fischer, E. R., Kochs, G., Grosse, R., García-Sastre, A., Vignuzzi, M., Johnson, J. R., Shokat, K. M., Swaney, D. L., Beltrao, P., and Krogan, N. J. (2020) The Global Phosphorylation Landscape of SARS-CoV-2 Infection. Cell. 182, 685-712.e19

